# Substructured population growth in the Ashkenazi Jews inferred with Approximate Bayesian Computation

**DOI:** 10.1101/467761

**Authors:** Ariella L. Gladstein, Michael F. Hammer

## Abstract

The Ashkenazi Jews (AJ) are a population isolate that have resided in Central Europe since at least the 10th century and share ancestry with both European and Middle Eastern populations. Between the 11th and 16th centuries, AJ expanded eastward leading to two culturally distinct communities, one in central Europe and one in eastern Europe. Our aim was to determine if there are genetically distinct AJ subpopulations that reflect the cultural groups, and if so, what demographic events contributed to the population differentiation. We used Approximate Bayesian Computation (ABC) to choose among models of AJ history and infer demographic parameter values, including divergence times, effective population size, and gene flow. For the ABC analysis we used allele frequency spectrum and identical by descent based statistics to capture information on a wide timescale. We also mitigated the effects of ascertainment bias when performing ABC on SNP array data by jointly modeling and inferring the SNP discovery. We found that the most likely model was population differentiation between the Eastern and Western AJ ~400 years ago. The differentiation between the Eastern and Western AJ could be attributed to more extreme population growth in the Eastern AJ (0.25 per generation) than the Western AJ (0.069 per generation).

## Introduction

We are at the cusp of an era that will utilize genomics to treat genetic diseases. As we enter the age of personal genomics, it is vital to fully understand the complex demographic factors that shape patterns of genetic variation. Without a better understanding of population dynamics, the promise of medical genetics cannot be fulfilled. Hidden population substructure, in which there are genetic differences between closely related populations can affect medical studies. Substructure resulting from different demographic histories can result in disparate frequencies of deleterious mutations. For example, rapid population growth causes an excess of rare variants, increasing the frequency of deleterious mutations in a population (1), while gene flow increases the frequency of common variants in a population. Knowledge of an individual’s ancestry and the demographic processes that contributed to their genomic architecture will help improve accuracy of medically relevant predictions based on genomic testing.

The Jewish communities from Western/Central and Eastern Europe, the Ashkenazi Jews (AJ), are an ideal population for studying the effects of complex demography, because they have a well documented history (2) with census records (3) and have been relatively isolated, with a high frequency of founder mutations associated with disease (4). The AJ have experienced an intricate demographic history, including bottlenecks, extreme growth, and gene flow, that affect the frequency of genetic diseases (see Gladstein and Hammer (5) for a more in-depth review of the literature and Carmi et al. (6), Xue et al. (7) for recent demographic inference). Different frequencies of mutations associated with diseases have been found among AJ communities (8), however most medical studies treat the AJ as one isolated population (see for example (4)). The historical record suggests there may have been a substantial difference in growth rates between the communities in Western/Central and Eastern Europe (3). Some authors suggest that the Eastern European AJ population size is unlikely to have reached observed 20th century levels without massive influx from non-Jewish Eastern Europeans (9). However, no genetic work has examined structured growth within the AJ and whether the recorded population growth in Eastern Europe is plausible without major external contributions. If left unresolved, this gap in our understanding of AJ population history could lead to deficient conclusions from medical studies.

The eastward expansion of the AJ settlement started after the 11th century and continued into the 16th century due to religious and ethnic persecution in Western and Central Europe (10). In the past few centuries, there have been two major culturally distinct communities of AJ - in Western/Central and Eastern Europe (11). This is reflected in the two primary Yiddish dialects - Western Yiddish and Eastern Yiddish, the latter of which can be subdivided into Northeastern (Lithuanian), Mideastern (Polish), and Southeastern (Ukrainian) (12) (Figure S1). Previous genetic studies have produced conflicting results on whether there is substructure within the AJ (8, 13–18), and is currently unclear whether major cultural subdivisions are reflected in genetic substructure of contemporary AJ. However, a recent study by Granot-Hershkovitz et al. (19) found substructure within AJ from SNP array data.

Our aim was to determine whether there are genetically distinct AJ subpopulations from Eastern and Western/Central Europe, and if so, whether differential population growth, gene flow, or both contributed to the population differentiation (Figure 1). We utilized genome-wide SNP data from 258 AJ with all four grandparents belonging to either western (n=19) or eastern (n=239) cultural groups and performed coalescent-based Approximate Bayesian Computation (ABC) to infer the most likely model for the AJ history. With current computational capabilities we were able to perform millions of genomic simulations at an unprecedented chromosome-size scale, allowing us to use segments identical by descent (IBD) for recent demographic inference, in addition to allele frequency based statistics, which are informative for the older history. The distribution of the length of IBD segments has the power to discern population structure, gene flow, and population size changes on a recent time scale (7, 20, 21). Previous methods using ABC used small loci, making chromosome-wide haploblock identification impossible, which is necessary for recent demographic inference. In addition, we used an ABC approach that takes into account SNP array ascertainment bias (22), so that SNP array data can be used without adverse effects of ascertainment bias. In this study we pushed the limit of computing capabilities by simulating whole chromosomes of hundreds of individuals, paving the way for future model-based inference of recent demographic history.

**Fig. 1.**
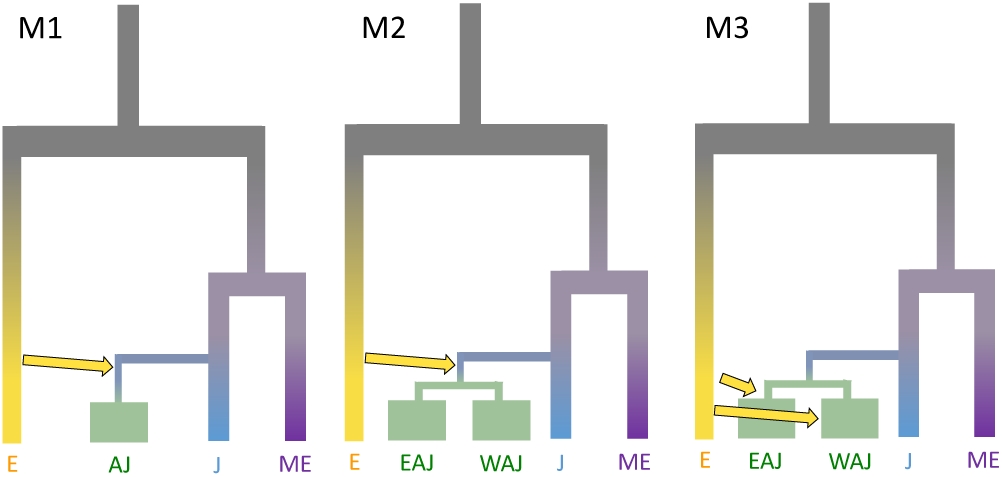
Three modeled demographic histories of the AJ. The first model (M1) has no substructure within the AJ. The second model (M2) has a population split between Eastern and Western AJ and one common admixture event from Europeans. The third model (M3) has a population split between Eastern and Western AJ and separate admixture events from the Europeans. Populations are labeled as E - European, AJ - Ashkenazi Jews, EAJ - Eastern Ashkenazi Jews, WAJ - Western Ashkenazi Jews, J - Sephardic/Mizrahi Jews, ME - Middle Eastern.

## Results

The PCA showed a similar pattern to other studies (19, 23, 24). However, there was also a separation of the Eastern and Western AJ, with the Western AJ further on PC2 from both the European and Middle Eastern clusters than the Eastern AJ (Figure 2). *Drop one in* PCA and ADMIXTURE revealed similar structure as PCA (Figures S6-S8).

**Fig. 2.**
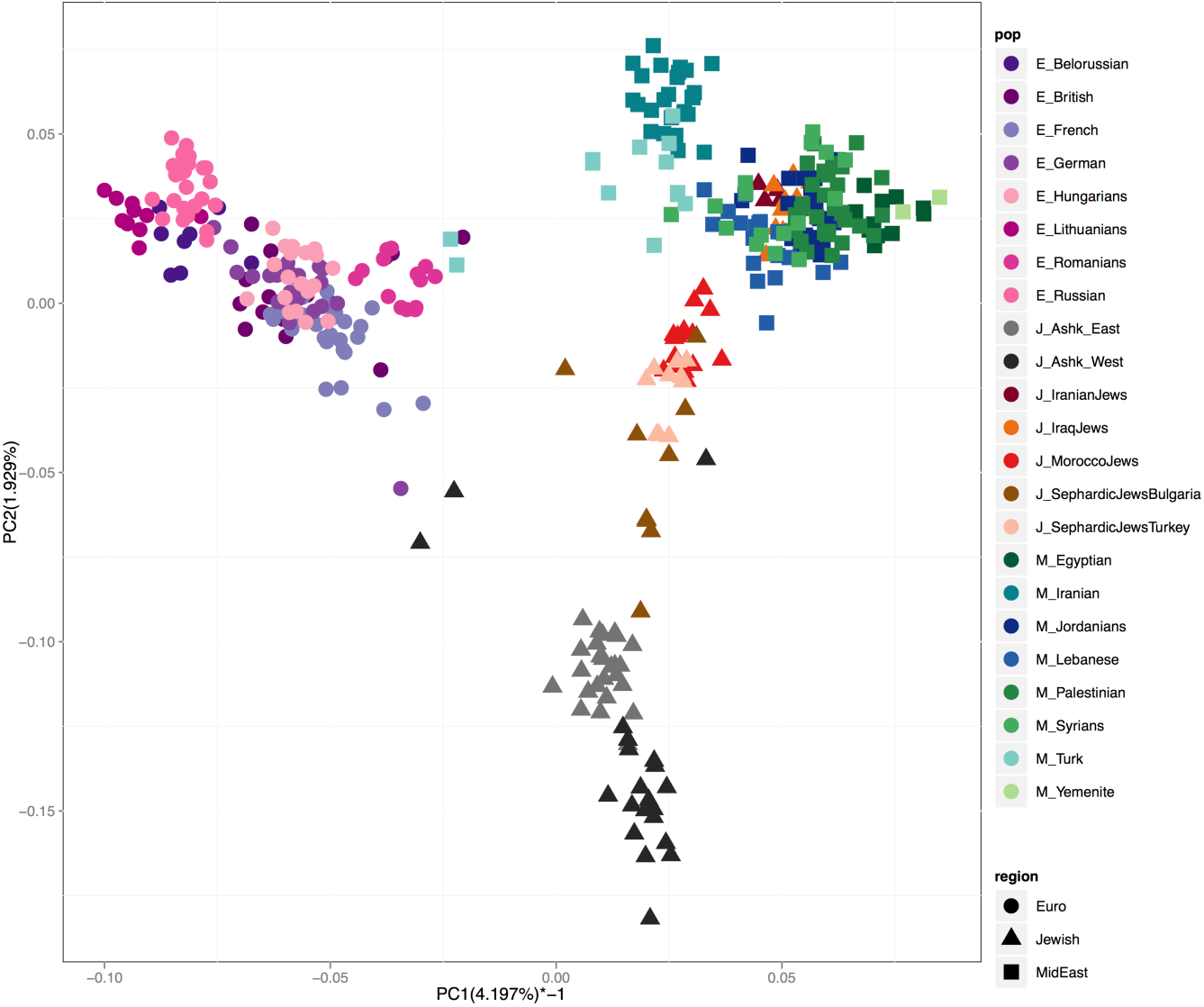
PC1 vs PC2 with Jewish, European, and Middle Eastern populations.

Eastern and Western AJ were significantly different in a variety of population genetic measures, all indicating that the Western AJ have a smaller effective population size or that the Eastern AJ have experienced greater gene flow from external populations (See supplemental section Test for substructure, figures S9-S15, and table S8).

## ABC

### Model choice

To test whether Eastern and Western AJ diverged into two subpopulations, and if so, whether different histories of population growth or gene flow contributed to the differentiation, we choose the best out of three demographic models (Figure 1) with ABC. Model 2 was the best fitting model with a Bayes factor of 2.01 and posterior probability of 0.67 (Table S9 and S10). Our observed Model 2 Bayes factor was greater than 92% of the Model 2 Bayes factors from the cross validations when Model 1 was the true model and greater than 86% of the Model 2 Bayes factors from the cross validations when Model 3 was the true model (Figure S18). Therefore, given that Model 2 was found to be the best model, it is unlikely that Model 1 or Model 3 is the true best fit.

### Parameter estimates

The estimations for the Middle Eastern, Jewish, and AJ populations are of particular interest to this study (Table 1, Figure 3). We used a generation time of 25 years to convert from generation to years. Since for the simulations we used a mutation rate of 2.5e-8, all parameter estimates were given in terms of that mutation rate. The parameter estimates are given in table 1. Parameter estimates from additional ABC analyses on the whole genome is given in supplemental tables S12 and S13.

**Fig. 3.**
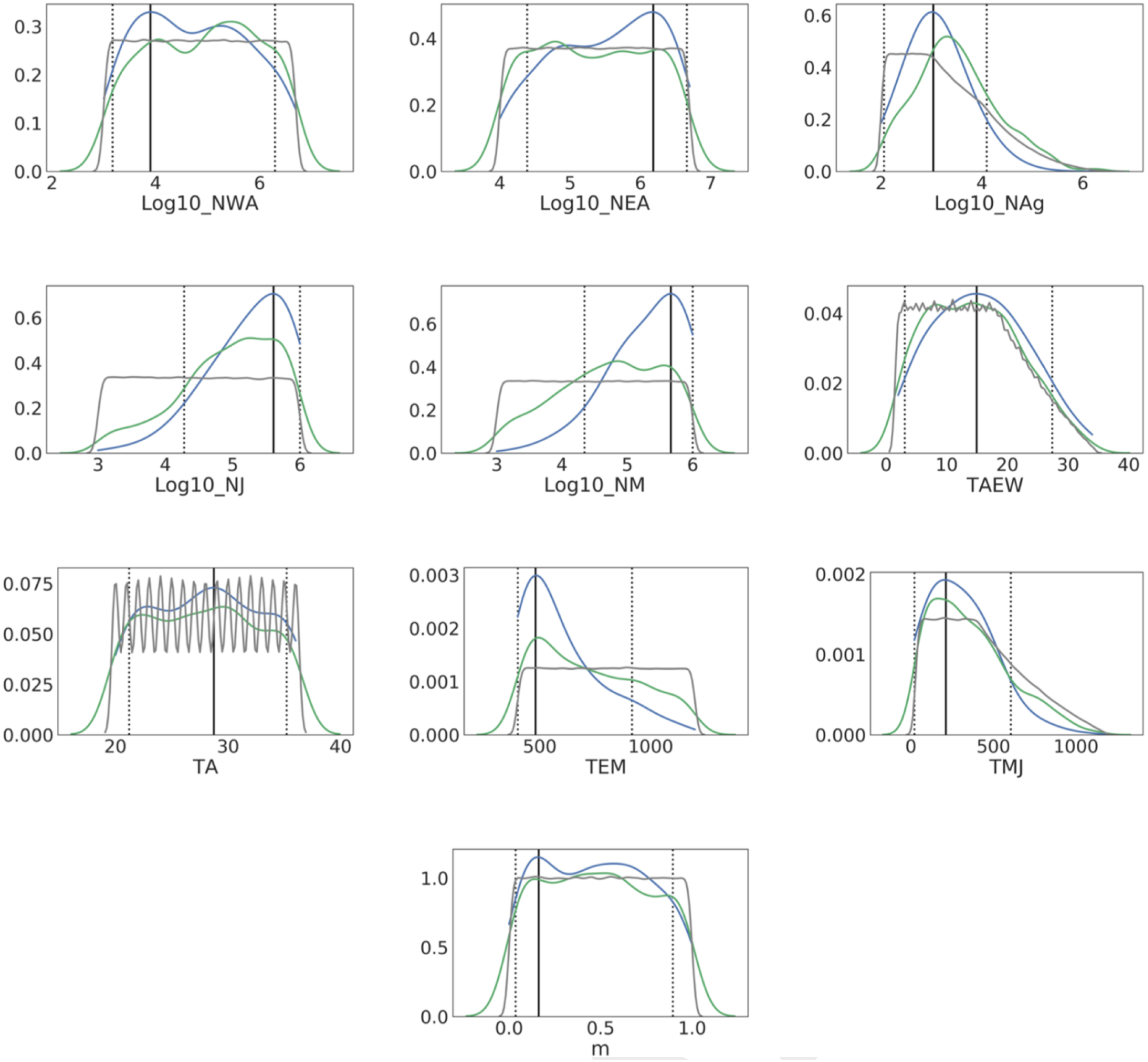
Posterior distributions of parameters of Model 2 from 1,446,125 simulations of chromosome 1. Black solid line is the mode of the posterior density and the dotted lines are the lower and upper limit of the 95 high posterior density interval. See Figure S3 for depiction of model parameters.

**Table 1.**
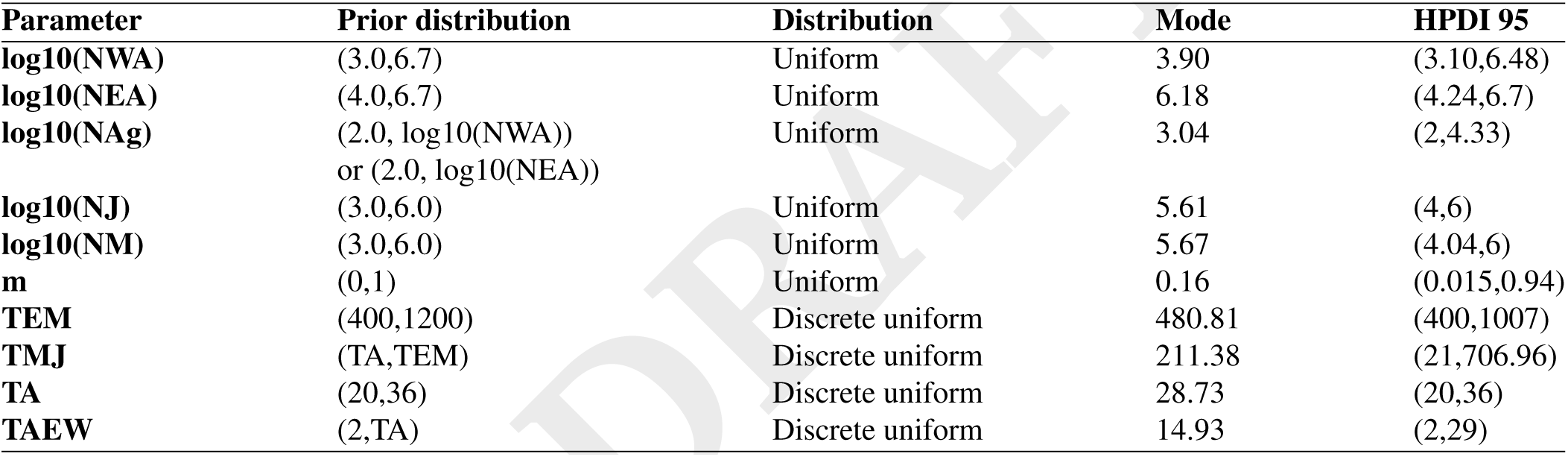
Prior distributions and parameter estimates of AJ demographic parameters Model 2 from 1,446,125 simulations of chromosome 1, with 10 PLS components and 1000 retained simulations. See Figure S3 for depiction of model parameters.

We found the probability that the Eastern AJ effective population was greater than the Western AJ effective population size from the joint posterior of the two parameters to be 0.69 (Figure 4). Given the estimated effective population sizes and divergence time from other Jewish populations, we estimated the exponential growth rate in Eastern and Western AJ to be 0.25 and 0.069 per generation, respectively.

**Fig. 4.**
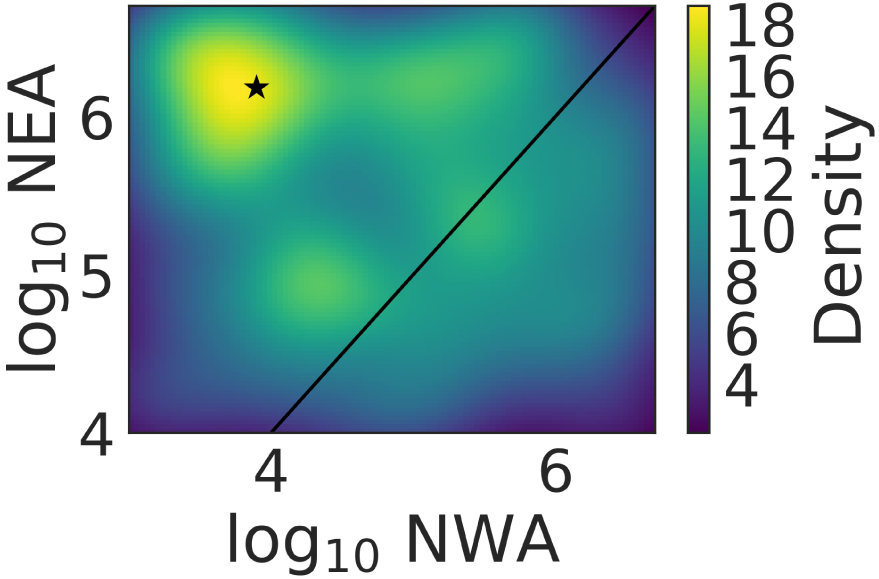
Joint posterior of effective population size in Eastern and Western AJ. The brighter colors indicate a higher posterior probability. The diagonal black line indicates where Ne in Western and Eastern AJ are equal. 69% of the joint density is greater in Eastern AJ than in Western AJ.

## Discussion

AJ traditionally trace their origin to the ancient Hebrews (Israelites) who lived as semi-herders in the Levant over 3000 years ago and have experienced a long subsequent history of migrations. The first known written accounts of “Israel”, in the central hill country of the southern Levant, is from the Merneptah Stele in 1207 BCE (25). The destruction of the First (587 BCE) and Second Temple (70 CE) contributed to the establishment of major Jewish centers outside of present-day Israel (26). While it is not clear what Jewish movement took place in Europe between the 1st and 4th centuries, migrations northward from Italy led to an established AJ community in the Rhine Valley by the 10th century (3). In the late Middle Ages and early modern period, Jews were expelled from much of Central Europe, and the Polish kings and nobility invited Jews to Poland (10).

From the end of the 16th century two major culturally distinct communities developed in Central and Eastern Europe (11), each speaking a different Yiddish dialect (12) and following different folk customs (e.g. dress, food, liturgical melodies, superstitions) (11). Throughout the Middle Ages in Central Europe, Jews were often expelled from their settlements and there were strict regulations on where Jews could live and what they could do to earn a living (11), while in Eastern Europe Jews could generally move freely and were protected by nobles, which fostered a sense of belonging (10). In the 19th century, Jews in Central Europe became more integrated in general life, with less focus on traditional and religious institutions (11), while Jews in Eastern Europe maintained Jewish learning and institutions and for the most part their lives revolved around Jewish religious tradition (10).

Our aim was to determine whether AJ with ancestry tracing to eastern and western cultural groups were genetically subdivided, and if so, what contributed to rapid population differentiation (Figure 1). Our models incorporated population structure, growth, and gene flow. Our first model represents the null hypothesis of cultural differentiation without genetic division between Eastern and Western AJ. This could be due to continuous gene flow between Eastern and Western AJ, insufficient time for genetic drift to cause differing allele frequencies, and/or similar rates of gene flow with Europeans in Eastern and Central Europe. Our second model includes a population split between Eastern and Western AJ allowing for different population sizes, following one common gene flow event with Europeans after the initial founder event. This could reflect low rates of gene flow between western and eastern AJ and/or different growth rates (with similar levels of gene flow from Europeans) after the western/eastern split. Our third model is similar to model 2; however, it allows for gene flow with Europeans separately in Eastern and Western AJ after the initial divergence of western and eastern AJ.

Our ABC analysis based on allele frequency spectrum and IBD statistics from extensive simulations of chromosome 1 rejected the null hypothesis of no substructure in the AJ (Model 1), and favored Model 2 over Model 3 (Table S9). While several factors could contribute to subdivision, we found that differential growth rates in the two subpopulations was a major factor. Our results support a model in which the AJ started from a small founder population upon initial arrival in Europe, followed by moderate population growth in Central Europe and massive population growth in Eastern Europe. Our conclusions from ABC are consistent with the observation of longer ROH and IBD and higher *F_IS_* in the Western AJ, which could reflect a smaller Ne than in Eastern AJ. The ADMIXTURE plot in Figure S9 is also consistent with more genetic drift in the Western AJ, which appears to have a higher proportion of an “AJ” component (i.e., other than two individuals that may be more recently admixed). Greater genetic drift may also be the explanation for the more extreme position of the WAJ in the PCA (Figure 2).

To compare our inferred growth estimates from ABC to census data, we found census data of Jews in Europe from the 12th to 20th centuries (3) (Figure S20, table S11). However, the source for the census data were not specific to AJ, and grouped Jews in Western and Southern Europe together, likely including Sephardi Jews from Southern Europe and the Balkans. Thus, we expect that the estimates of the population sizes for the Jews of Western and Southern Europe is an overestimate of the population sizes of Western AJ. We fit an exponential model to the census data from the years 1170 to 1900. We found the same order of magnitude in Eastern AJ and high concordance in Western AJ for our estimates of growth rates based on genomic (0.25 per generation in Eastern AJ and 0.069 per generation in Western AJ) and census data (0.18 ± 0.017 per generation in Eastern European Jews and 0.081 ± 0.026 per generation in Western/Southern European Jews). Coventry et al. (27) estimated similar growth rate in Europeans from census data (0.115 per generation since 1600) (28), and a 10-fold higher growth rate in Europeans from genetic data (1.094 per generation).

The estimated census population growth in the Eastern AJ has been referred to as the “Demographic Miracle” (2), and has been the center of much debate with some arguing that it is impossible biologically, without massive conversion or intermarriage (9). However, from our ABC analysis, since Model 2 was favored over Model 3, we did not find that gene flow from Europeans was necessary to increase the effective population size in the Eastern AJ relative to Western AJ. Our genetic results corroborate DellaPergola (3)’s conclusion based off of demographic analysis of census data that the rapid growth of Eastern AJ is feasible without mass immigration or large-scale conversions to Judaism.

A potential, large demographic contribution to EAJ through conversion/gene flow has been hypothesized to come from the Khazars (Figure S22). Historical documents suggest that the Khazars were a Turkic people, with an empire that spanned parts of modern day Ukraine, southern Russia, and the Caucasus from the 7th to 11th century (Brook 2006). While some documents suggest that Khazarian royalty converted to Judaism for political reasons, it is unknown how much of the Khazarian population converted (29). Elhaik (30) and Behar et al. (31) have both addressed the possibility of substantial Khazarian contribution into the unstructured AJ gene-pool, with Elhaik (30) in favor of the Khazarian hypothesis and Behar et al. (31) opposed. While we did not explicitly test a model of Turkic gene flow to the Eastern AJ in our ABC analysis, our results indicate that the Eastern AJ could have undergone the observed growth without contribution from outside populations, such as the Khazars. In addition, we found with an ad-hoc resampling simulation, with Chuvash as a proxy for the Khazars and Western AJ as a proxy for ancestral Eastern AJ, that adding Chuvash gene flow to the Western AJ did not create an admixed population similar to Eastern AJ (Figure S23). Meaning that Eastern AJ are unlikely to be an admixed population with Turkic and Western AJ sources. Furthermore, if the Khazars had contributed substantially to the population size of AJ in Eastern Europe, we would expect a population size increase around the time of the fall of the Khazarian Empire, or shortly after. However, according to the census data Eastern AJ experienced high growth rates relative to the Western/Southern Jews in the 16th and 20th centuries, well after the fall of Khazaria.

If the increased population growth in the Eastern AJ was not due to contributions from outside populations, there must be other reasons that account for the higher growth rate in the Eastern AJ over the Western AJ. The primary historical explanation for the AJ population increase in Eastern Europe was better political, economic, and social conditions (11). In Central Europe, discriminatory legislation of the late Middle Ages and early modern period meant that very few Jews lived there (11). There were often legal limitations on the number of Jewish families - irrespective of where they lived (11). In the 19th century cramped ghettos and integration into non-Jewish society contributed to low birthrates (32). Unlike in other regions of Europe, Jewish communities in Eastern Europe did not have limitations from the government on the number of Jewish marriages permitted, and did not have strict residence restrictions (32). Adherence to religious and traditional norms and the economic structures encouraged early marriage and high fertility in Eastern Europe (32). Additionally, generally low mortality rates among the AJ, particularly low infant mortality, contributed to the high growth rate. These factors could have contributed to the high population growth in the Eastern AJ, leading to rapid genetic differentiation between the Eastern and Western AJ.

### Implication for genetic diseases

The AJ have an unusually high prevalence of more than 50 known disease-associated mutations (33), primarily because of founder effects resulting from population bottlenecks (8, 34). Both Risch et al. (8) and Slatkin (34) found evidence of founder events ~11 and ~5 centuries ago. Slatkin (34) tested for neutrality and founder effect in several disease-associated alleles found predominantly in AJ. He used LD with a linked marker allele and a historical demographic model, assuming no subdivision in the AJ. Additionally, Risch et al. (8) examined the allele-frequency distributions and estimated coalescence times of several disease-causing mutations at increased frequency in the AJ and found that the more recent founder mutations were restricted to Lithuanians, and the older founder mutations were present across all AJ subpopulations. While different histories of growth in a substructured population could have implications on the frequency of disease, no previous work has modeled population structure and population size changes in the AJ.

The different growth rates in Western and Eastern AJ may have varying effects on the frequency of deleterious mutations in the subpopulations. Extreme recent population growth in the Eastern AJ increases the likelihood of new mutations on a background of high homozygosity because of their extreme founder event. Although we do not observe this in our SNP array data, it is unlikely to be observable from standard SNP arrays. Extreme recent population growth means that individuals of Eastern AJ descent may have an increased probability of being homozygous for deleterious alleles. The Western AJ are less likely than the Eastern AJ to have homozygous segments that contain pathogenic alleles. While the Western AJ have a higher probability of a site being homozygous, they did not have as extreme population growth as the Eastern AJ, and thus are less likely to have new mutations leading to pathogenic alleles. It has previously been inferred that numerous deleterious mutations at high frequency in the AJ occurred when the Lithuanian Jewish community was founded and expanded, which according to census data, was one of the sources of extreme population growth (8, 35). It would be important for AJ individuals to be aware of the effect the extreme growth rate has on genetic disorders in Eastern AJ. However, today there are few AJ with ancestry only from Western/Central Europe, because of the population growth in the Eastern AJ and mixing between Jews outside of Europe. This study serves as an example to demonstrate that recent population differentiation is possible and may have genetic consequences that impact the health of the individuals today.

### In context of other studies

Since we used a combination of allele frequency spectrum and identity by descent based statistics, the old demographic events (TEM and TMJ) could be effected by the mutation rate, while recent events should be mostly effected by recombination rate. Therefore the mutation rate used in the simulation should not effect the majority of the inferred parameters. However, the parameter estimates can be adjusted to the 1.2e-8 (36) or 1.44e-8 (37) mutation rate by doubling the estimated divergence times and effective population sizes.

Our estimate of the Middle Eastern and European divergence time corresponds with other published estimates (6, 38). Our estimate of the divergence time of the Ashkenazi Jews from the non-Ashkenazi Jews overlaps with estimates of when the Ashkenazi Jews experienced a population size reduction (6). Our estimate of current effective population size in the Eastern AJ is in agreement with Carmi et al. (6)’s estimate of AJ effective population size and our estimate of the founder effect size is slightly larger, but the 95 high posterior density interval overlaps (6).

Previous genomic scale demographic studies did not take into account population substructure in their sampling strategies (6, 7, 21, 39). While our conclusion of substructure in the AJ is consistent with results from studies using haploid loci and classical markers (8, 13–17), it is contrary to Guha et al. (18) conclusion that the AJ are not genetically subdivided. However, Guha et al. (18)’s PCA results of AJ with ancestry from different countries does not provide support for or against our conclusion of subdivision between the Eastern and Western AJ, because they do not use historically/culturally motivated AJ groupings.

Xue et al. (7) inferred an admixture event with Southern Europeans around the time of the AJ founder event ~24-49 generations ago, and at least one admixture event ~10-20 generations ago; however, they were not able to infer whether this more recent event involved western or eastern non-Jewish Europeans. While the uncertainty could be do to limitations in their methods to distinguish between western or eastern European sources, it is plausible that their uncertainty was due to their use of the AJ as a single population. Potentially, if they had performed their analyses separately with Eastern AJ and Western AJ, they would have been able to more clearly identify the sources of the later admixture events.

### Caveats and Future directions

We attempted to keep the model as simple as possible, while still providing insight into substructure in the AJ. For this reason, we did not include European gene flow to Sephardic Jews, which likely pushed back the divergence time between the Sephardic Jews and Middle Eastern population, due to the European lineages in the Sephardic Jews. This may also have contributed to a recent estimate of divergence time between the AJ and Sephardic populations and contributed to an estimated large effective population size in the Sephardic Jews. Additionally, we did not simulate gene flow between Eastern and Western AJ because we believed that we would not have enough power to infer gene flow between Eastern and Western AJ. It is possible there was continuous gene flow between the Eastern and Western AJ, which could cause our estimated divergence time to be more recent. Additionally, while model 2 explicitly includes growth after EAJ diverged from WAJ, we did not explicitly test whether growth started before or after divergence.

While we attempted a-priori to pick summary statistics that would capture the old and recent history, it is possible that our set of summary statistics were not informative for all aspects of the model. In particular, for Model 2 we inferred a surprisingly low proportion of European gene flow into the AJ. In comparison, recent studies have estimated approximately 50% admixture from Europeans (6, 7). We believe that our combination of summary statistics may not have been informative for European gene flow, and we are not confident in the admixture proportion estimates in this study.

Regarding AJ demographic history we would like future studies to address three topics - more detail regarding population growth, the source of small effective population size, and more complex admixture history. Future studies should compare models of instantaneous, exponential, and logistic growth. Future studies could also distinguish between a drift caused by a small effective population size in a randomly mating population versus drift caused by a small effective population size due to preferential mating of more closely related individuals. Finally, future studies could use more sensitive local ancestry estimation in the Eastern and Western AJ with multiple European sources and multiple admixture events, with distinction between multiple events and continuous admixture.

## Materials and Methods

### Dataset

High density SNP array in Jewish, European, and Middle Eastern populations, and whole genome sequence data in world-wide populations were used (Tables S1 and S2). An additional 231 AJ samples were genotyped on Illumina Omni Express chip for 730K markers. AJ samples were collected in the United States and Israel and information on the grandparental country of origin was provided, with a total of 13 European countries. We defined the AJ as Eastern or Western based on their grandparental country of origin, according to Yiddish dialect borders (Figure S1). All included samples had four AJ grandparents from Eastern or Central Europe (we did not include AJ individuals of mixed Eastern and Central European ancestry).

### Approximate Bayesian Computation

We used Approximate Bayesian Computation (ABC) to choose the best supported demographic model and to obtain the posterior distribution of model parameters values under the best-supported model. We used the ABC-GLM (General Linear Model) approach introduced by Leuenberger and Wegmann (40) and implemented in the program `ABCtoolbox` 2.0 (41). An overview of the ABC pipeline is shown in figure 5. The following sections provide more details on each of the ABC steps (reproducible step-by-step instructions are available on Bitbucket, https://bitbucket.org/agladstein/macsswig_simsaj).

**Fig. 5.**
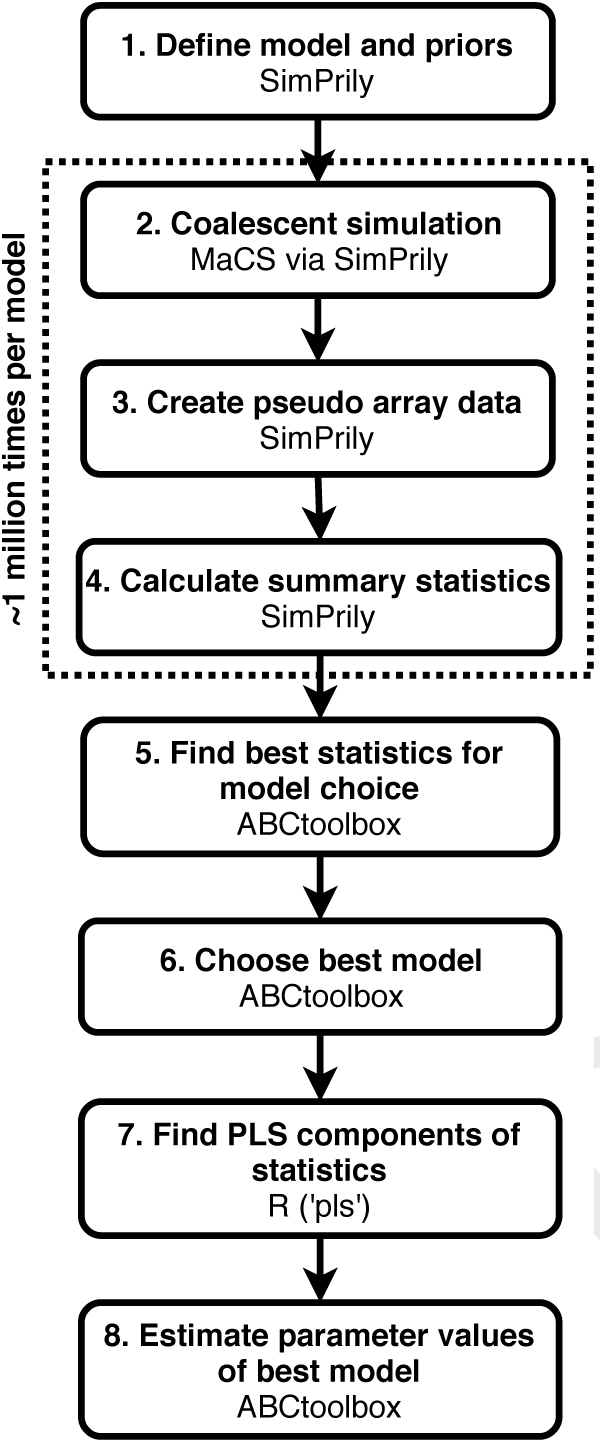
Overview of ABC pipeline. The eight main steps of the pipeline with the tools used to perform them (41–44).

### Demographic models

We simulated three models of AJ demographic history, with an incorporated SNP array ascertainment scheme based on the three HapMap populations, CEU, CHB, and YRI (22) (Figures S2, S3, S4). The priors were based on previously inferred parameter values (6, 22, 45) and documented history (2) (Table S3). Based on the sample sizes from the observed data, we simulated 9 YRI, 9 CEU, 4 CHB, 38 AJ (or 19 Eastern and 19 Western for models 2 and 3), 14 Jewish, and 14 Middle Eastern. We reduced the simulation sample size of the AJ (specifically Eastern), Jewish, and Middle Eastern populations to reduce the run-time of the simulations. The three models are the same until the AJ diverge from the other Jewish populations. Model 1 represents the null hypothesis of no population structure in the AJ, while in model 2 and model 3 there is a population split in the AJ. In model 2, gene flow from Europeans to the AJ occurs before the AJ split. In model 3, gene flow from Europeans occurs after the split in the AJ, allowing for different proportions and times of gene flow in Eastern and Western AJ. Both Model 2 and Model 3 allow for independent population growth in the Eastern and Western AJ. The three models are described in detail in tables S4, S5, S6, with their appropriate MaCS (43) commands.

### Simulations

We performed about 1 millions simulations of chromosome 1 and calculated summary statistics with SimPrily-alpha (42), which uses a modified version of Markovian Coalescent Simulator (MaCS) (43). We ran the simulations in parallel on high throughput clusters (46–51) (figure S5). We used a mutation rate of *μ*=2.5e-8 (52) and a HapMap recombination map (53), with the recombination rate, *ρ*=1e-8.

We made pseudo SNP arrays from the simulated whole chromosome 1 based on real SNP arrays and a discovery sample. We followed Quinto-Cortés et al. (22)’s method, and used random samples from YRI, CEU, and CHB as the SNP discovery set (prior of 2,20 chromosomes), and used a random derived allele threshold (0.05,0.1) to find SNPs to use for the pseudo array. To create the pseudo array we found the closest simulated sites to the real array that had a minor allele frequency greater than the threshold in the discovery sample. By taking into account ascertainment bias in our simulated model were able to recover rare alleles not present in the SNP array data (Figure S17).

### Summary statistics choice

We included allele frequency spectrum based summary statistics to capture information on the older parts of the demography and included IBD statistics to summarize information on recent history (Table S7). We used a total of 181 summary statistics, consisting of the number of segregating sites, number of singletons, number of doubletons, Tajima’s D,*F_ST_*, mean and median length, variance of length, and number of shared IBD greater than 3Mb and 30Mb.

To choose the best model, we found the best sets of summary statistics for model choice from a pruned subset of summary statistics, using the greedy search algorithm in `ABCtoolbox` (see Supplemental methods). In order to avoid correlations among statistics and to preserve the informativeness of the data, we transformed our statistics with Partial Least Squares (PLS) and used the first 10 PLS components for the parameter estimation.

### Model Choice and parameter estimation

For both model choice and parameter estimation, in order to maximize our power we used a reduced set of 9, 11, and 13 parameters for models 1, 2, and 3, leaving the remaining parameters as free parameters, which were not inferred as part of ABC. We compared the three models using 1,275,807 simulations with `ABCtoolbox`, which chooses the best model by calculating Bayes factors for each compared model. The Bayes factors are calculated from the marginal densities of the models (41). We then inferred the parameters of the best model with 1,446,124 simulations using `ABCtoolbox`, which applies a General Linear Model regression adjustment to estimate the posterior densities of parameters (41). We used the mode of the posterior densities as the parameter estimates.

### Validation of model choice and parameter estimation

We used cross-validation with `ABCtoolbox` to assess the accuracy of model choice and parameter estimates of the best model. Cross-validation executes the model choice and parameter estimation process 1000 times by choosing one of the simulated data sets as the observed data. From this, we obtained a confusion matrix with estimates of false positives and false negatives. Due to the way the cross validation works, it is necessary for all models to have the same summary statistics. Therefore, we forced model 1 to have the same summary statistics as model 2 and 3 by randomly splitting the simulated AJ into two groups and calling one Eastern and one Western.

## ACKNOWLEDGEMENTS

An allocation of computer time from the UA Research Computing High Performance Computing (HPC) at the University of Arizona is gratefully acknowledged.

We would like to thank Mats Rynge for his invaluable help setting up the Pegasus workflow and running it on the Open Science Grid. This research used resources provided by the Open Science Grid, which is supported by the NSF award 1148698, and the U.S. Department of Energy’s Office of Science.

This material is based upon work supported by the National Science Foundation under Award Numbers DBI-0735191 and DBI-1265383 and we thank CyVerse for their External Collaborative Support program.

This work used the Extreme Science and Engineering Discovery Environment (XSEDE), which is supported by National Science Foundation grant number ACI-1548562. The computations were conducted on the Comet supercomputer, which is supported by NSF award number ACI-1341698, at the San Diego Supercomputing Center (SDSC), the Jetstream cloud environment, which is supported by National Science Foundation award number NSF-1445604, at Indiana University and Texas Advanced Computing Center, and the Bridges supercomputer, which is supported by NSF award number ACI-1445606, at the Pittsburgh Supercomputing Center (PSC).

This research was performed using the compute resources and assistance of the UW-Madison Center For High Throughput Computing (CHTC) in the Department of Computer Sciences.

Pegasus is funded by the US NSF under OAC SI2-SSI program, grant #1664162.

